# Motile epipelic diatoms exhibit dynamic aggregation behaviour altered under oxidative perturbation

**DOI:** 10.64898/2026.09.03.749072

**Authors:** Alexandre Desparmet, Cédric Hubas

## Abstract

Epipelic diatoms continuously adjust their position within sediments, yet how individual motility contributes to the spatial organization and temporal dynamics of collective aggregation and how this behaviour responds to environmental conditions remain poorly understood. We resuspended a natural assemblage dominated by *Pleurosigma strigosum* and *Gyrosigma balticum* in filtered seawater and monitored aggregation dynamics over 72 h under a dark control condition or under 24 h exposure to a nominal initial H_2_O_2_ concentration of 400 µmol L^−1^, followed by a 48 h post-exposure period. Time-lapse imaging and quantitative image analysis revealed rapid aggregation and continuous remodeling into dynamic, spatially complex multicellular networks. In the control, aggregate number declined from 11,767 to 3,237 and surface coverage from 24.4% to 10.3%, with pronounced temporal fluctuations. Under H_2_O_2_, aggregation followed a distinct, less variable trajectory, with fewer aggregates and persistent differences following the cessation of H_2_O_2_ exposure. These observations suggest that dynamic aggregation behaviour may contribute to biofilm organization and resilience, while not distinguishing active regulation from emergent cell–cell–matrix interactions.

---

The surface of cohesive muddy sediments in intertidal mudflats hosts highly productive microphytobenthic communities embedded within extracellular polymeric substance (EPS)-rich matrix (Hubas et al., 2018; Hubas et al., 2023; Underwood et al., 2022). These biofilms, often dominated by motile epipelic diatoms, contribute substantially to coastal ecosystem services through primary production, biogeochemical cycling, support of food webs, and sediment stabilization (Passarelli et al., 2018; Pinckney, 2018; Underwood et al., 2026). Epipelic diatoms experience rapid fluctuations in irradiance, temperature, desiccation, pH, salinity, nutrient availability, and other environmental conditions, some of which can promote the production and accumulation of reactive oxygen species (Waring et al., 2010; Foyer and Hanke, 2022; Morris et al., 2022). Their motility allows individual cells to reposition rapidly within sediments in response to environmental variation and thereby regulate their exposure to potentially stressful conditions (Consalvey et al., 2004; Jesus et al., 2023; Desparmet et al., 2026). This gliding motility is driven by an actin-myosin-based cytoskeletal system coupled to the secretion of extracellular mucilage fibrils through the raphe, enabling cell adhesion and traction on the substratum at low metabolic cost (Poulsen et al., 1999; Marques Da Silva et al., 2020).

Although the motile responses of individual epipelic diatoms are increasingly well characterized, how individual motility contributes to the spatial organization and temporal dynamics of collective aggregation remains poorly understood. Aggregation, defined here as the transition of initially dispersed cells toward multicellular associations, is well established in diatoms and other microalgae and may involve EPS-mediated adhesion and other cell–cell and cell–matrix interactions (Wang et al., 2021; De Carpentier et al., 2022; Lambert et al., 2026). However, little is known about how aggregation emerges and how this collective organization evolves over time in natural epipelic assemblages. It also remains unclear whether reactive oxygen species may influence this dynamic and to what extent the resulting patterns reflect biologically regulated behaviour rather than emergent consequences of motility and cell–cell–matrix interactions. Here, we characterized the aggregation dynamics of a natural assemblage of motile epipelic diatoms following resuspension and examined how their temporal dynamics differed during H_2_O_2_ exposure and the subsequent postexposure period.

A natural microphytobenthic community dominated by epipelic diatoms, particularly *Pleurosigma strigosum* and *Gyrosigma balticum* (Fig. 1E), and containing a small meiofaunal component (e.g. nematodes, copepods, and ostracods), was collected from fine-grained muddy sediment in a disused outdoor breeding pond subject to a semi-diurnal tidal regime at the Concarneau Marine Station, France (26 July 2026, 10:00; 47°52ʹ05.8502BAN, 3°55ʹ00.51ʺW). Samples were allowed to settle for 24 h at 22°C under ambient laboratory light without direct illumination. Two liters of microalgal suspension were prepared by gently removing the cohesive, EPS-rich upper ∼1 mm of microphytobenthos with tweezers and resuspending the material by agitation in filtered site seawater, as described in Desparmet et al. (2026).

**Figure 1.**
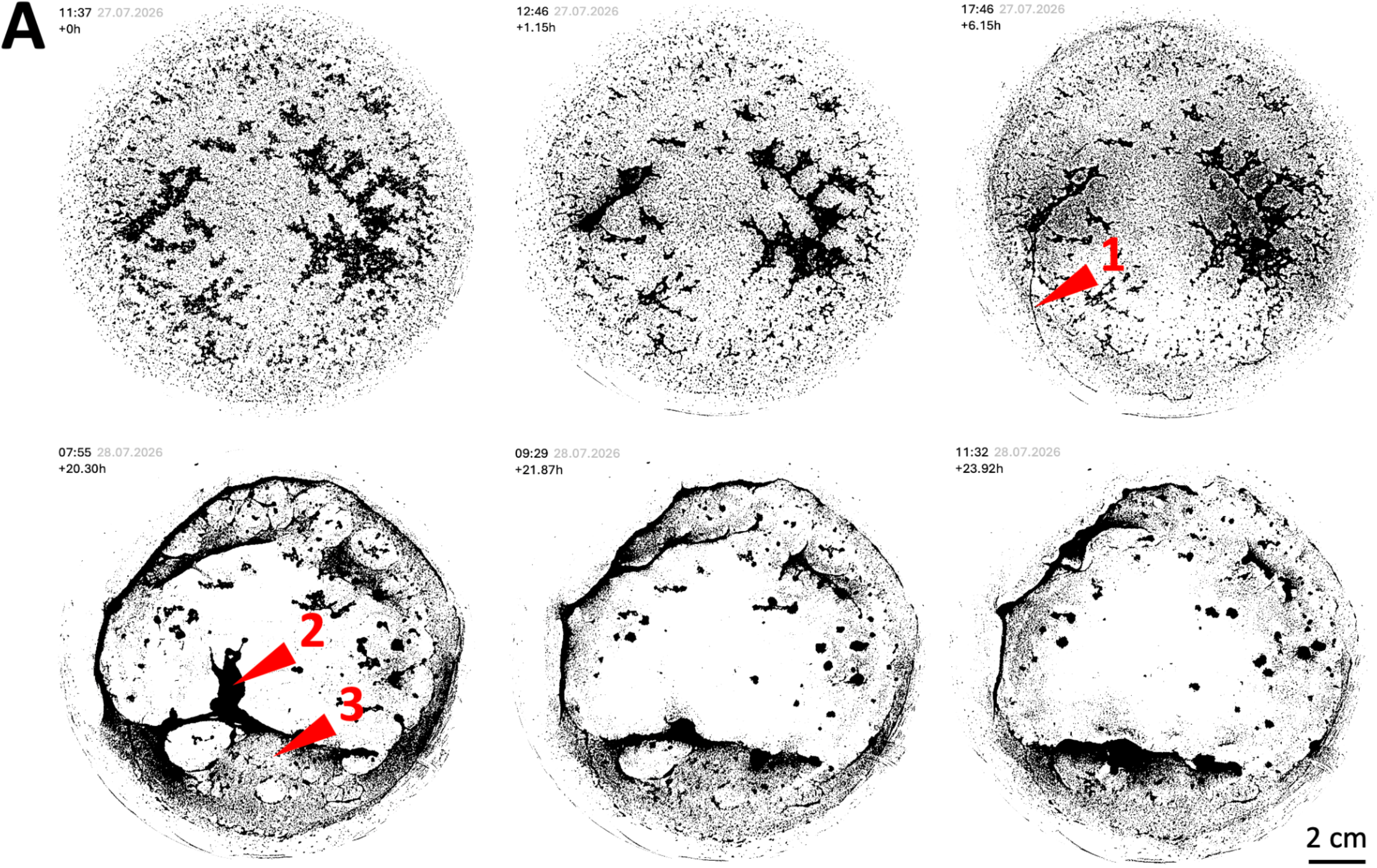
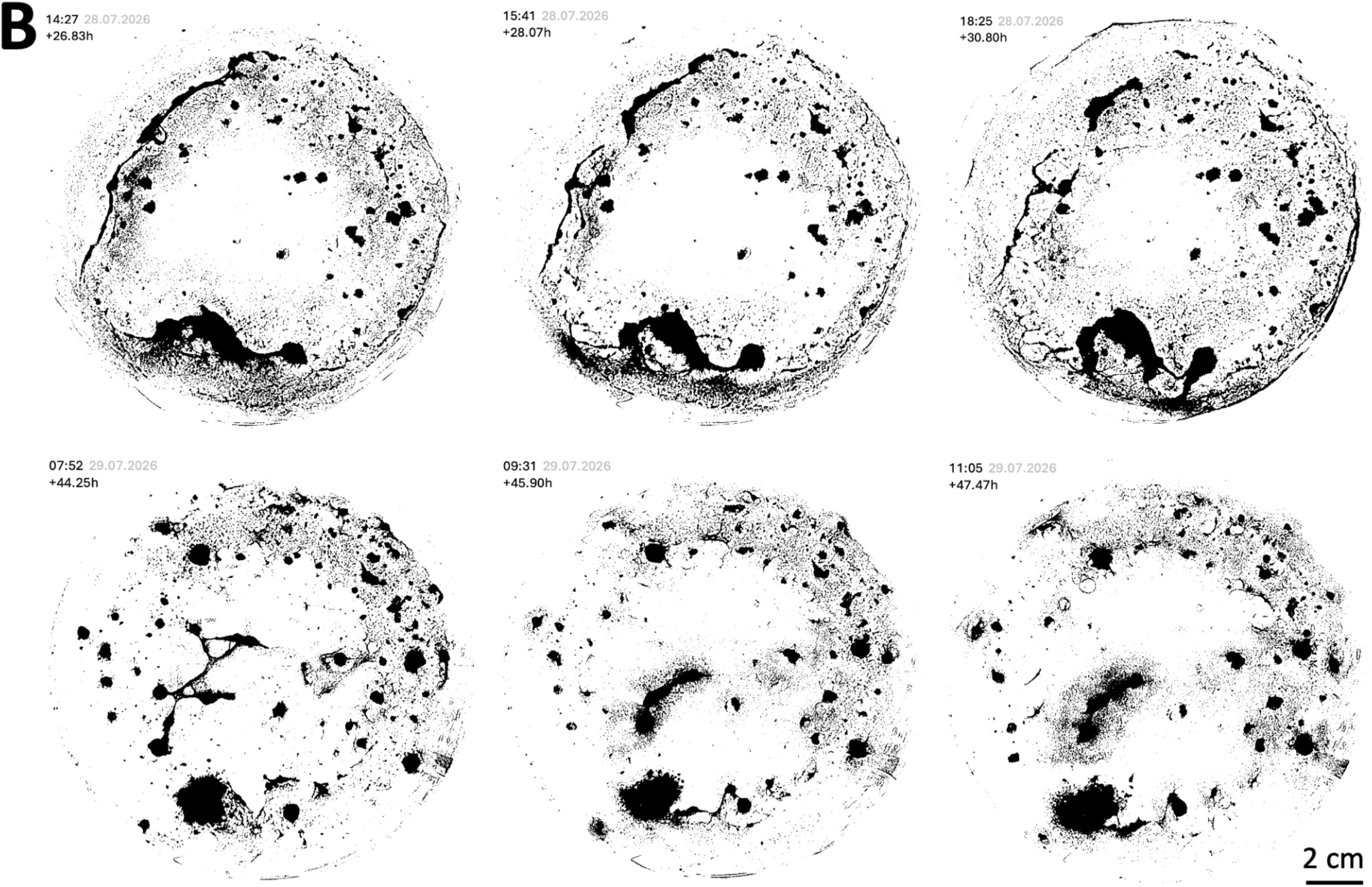
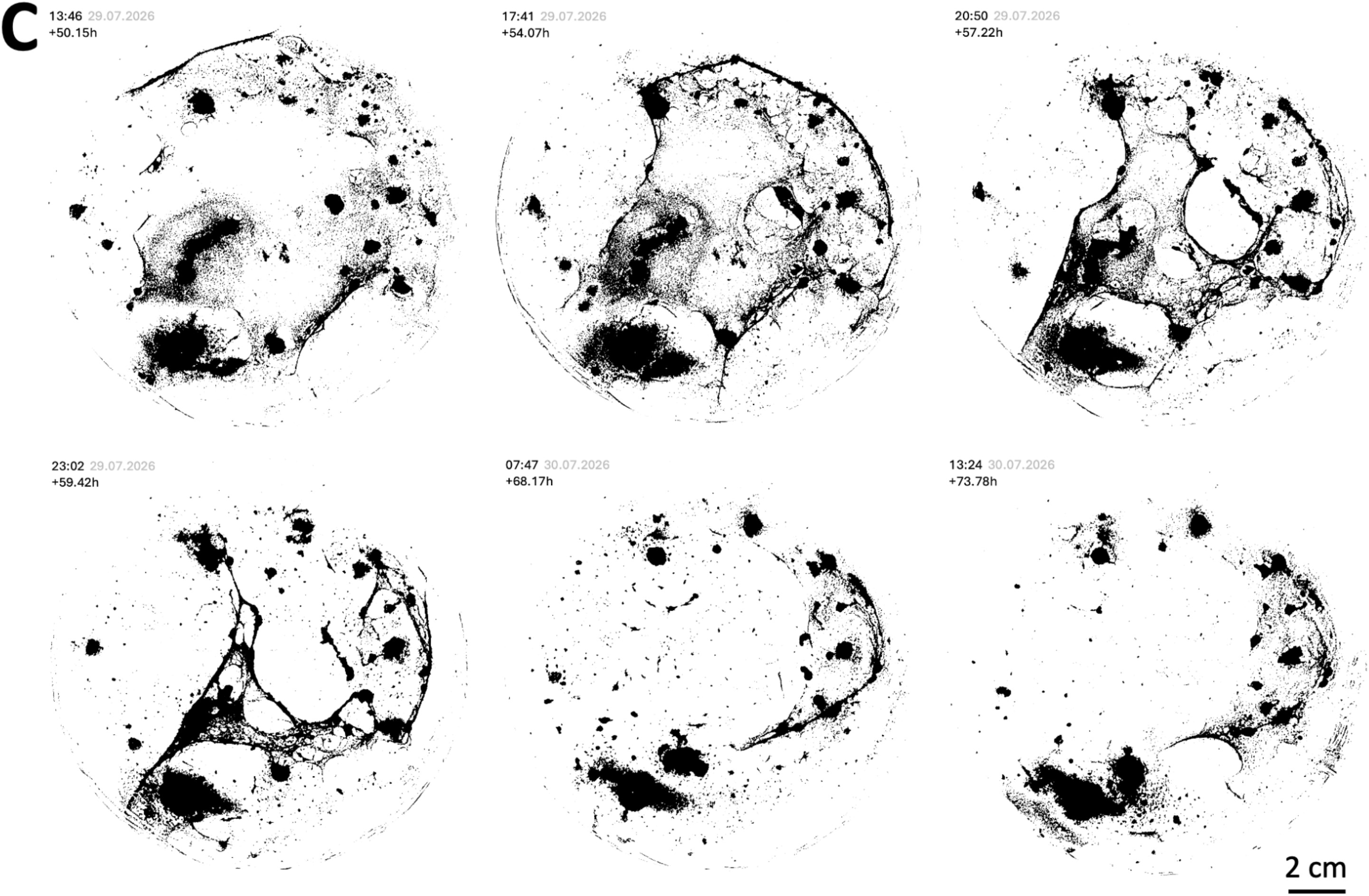
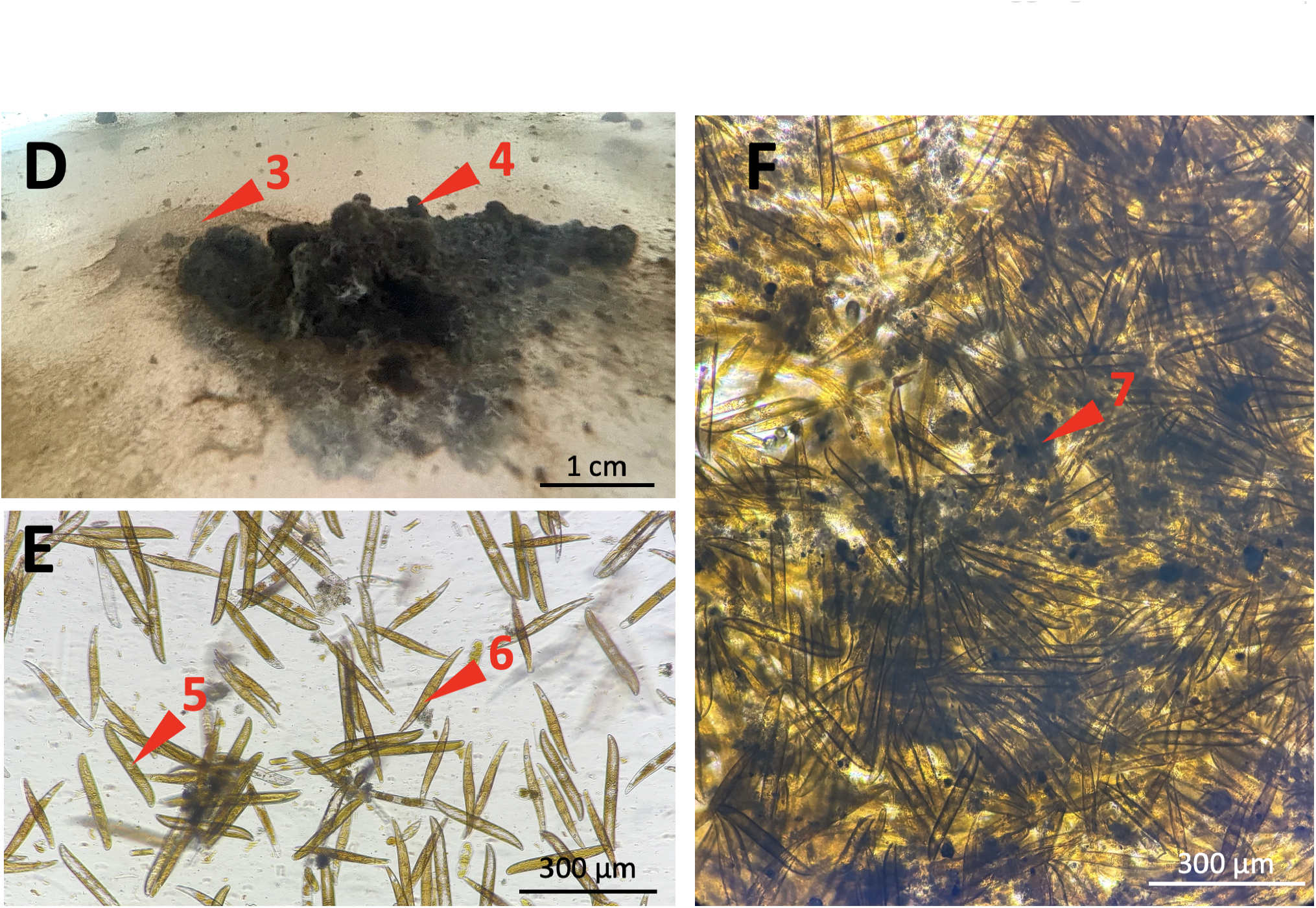
Temporal dynamics and structural organization of diatom aggregation under the control condition. Representative images of the natural microphytobenthic assemblage monitored over 72 h following resuspension from sediment. Images were repeatedly acquired from the bottom of the same 3-L Erlenmeyer flask containing 1 L of suspension. **A–C**. Representative temporal sequences showing changes in aggregation patterns during 0–24 h (A), 24–48 h (B), and 48–72 h (C). Arrows in panel A indicate examples of a filamentous structure^**(1)**^, a compact aggregate^**(2)**^ and a biofilm-like mat^**(3)**^; **D**. Macroscopic view of an aggregate showing its spatially complex morphology, including mushroomlike macrocolony structures^**(4)**^ (as described in bacterial biofilms; Flemming et al. 2011); **E**. Representative lightmicroscopy image of the natural assemblage, dominated by motile epipelic diatoms including *Gyrosigma balticum*^**(5)**^ and *Pleurosigma strigosum*^**(6)**^; **F**. Light-microscopy image of a dense aggregate showing closely packed diatom cells. The darker appearance of densely aggregated regions^**(7)**^ reflects high optical density, likely resulting from local EPS accumulation. Quantitative image analysis of aggregate number, surface coverage, and their temporal dynamics is provided in Supplementary Fig. 1 and Supplementary Table 1.

The suspension, adjusted to ∼5 x10^6^ diatoms L^−^1 as assessed by microscopic cell counting, was divided between two 3-L borosilicate Erlenmeyer flasks (Pyrex), each containing 1 L. One flask served as the control condition (n = 1), while 42 µL of stabilized H_2_O_2_ stock solution (9.7912 mol L^−1^; Sigma-Aldrich) was added to the second flask, corresponding to a nominal initial H_2_O_2_ concentration of ∼400 µmol L^−1^ (n = 1). After 24 h of H_2_O_2_ exposure, the H_2_O_2_-containing seawater was removed while retaining the microalgal material in the flask and replaced with filtered site seawater without added H_2_O_2_ for the remaining 48 h, corresponding to the post-exposure period. Both flasks were maintained at 22°C in darkness (<1 µmol photons m^−^2 s^−^1) and changes in community spatial organization and aggregation dynamics were monitored by repeated imaging from the bottom of each flask at several time points over the 72 h experiment (Figs. 1, 2).

**Figure 2.**
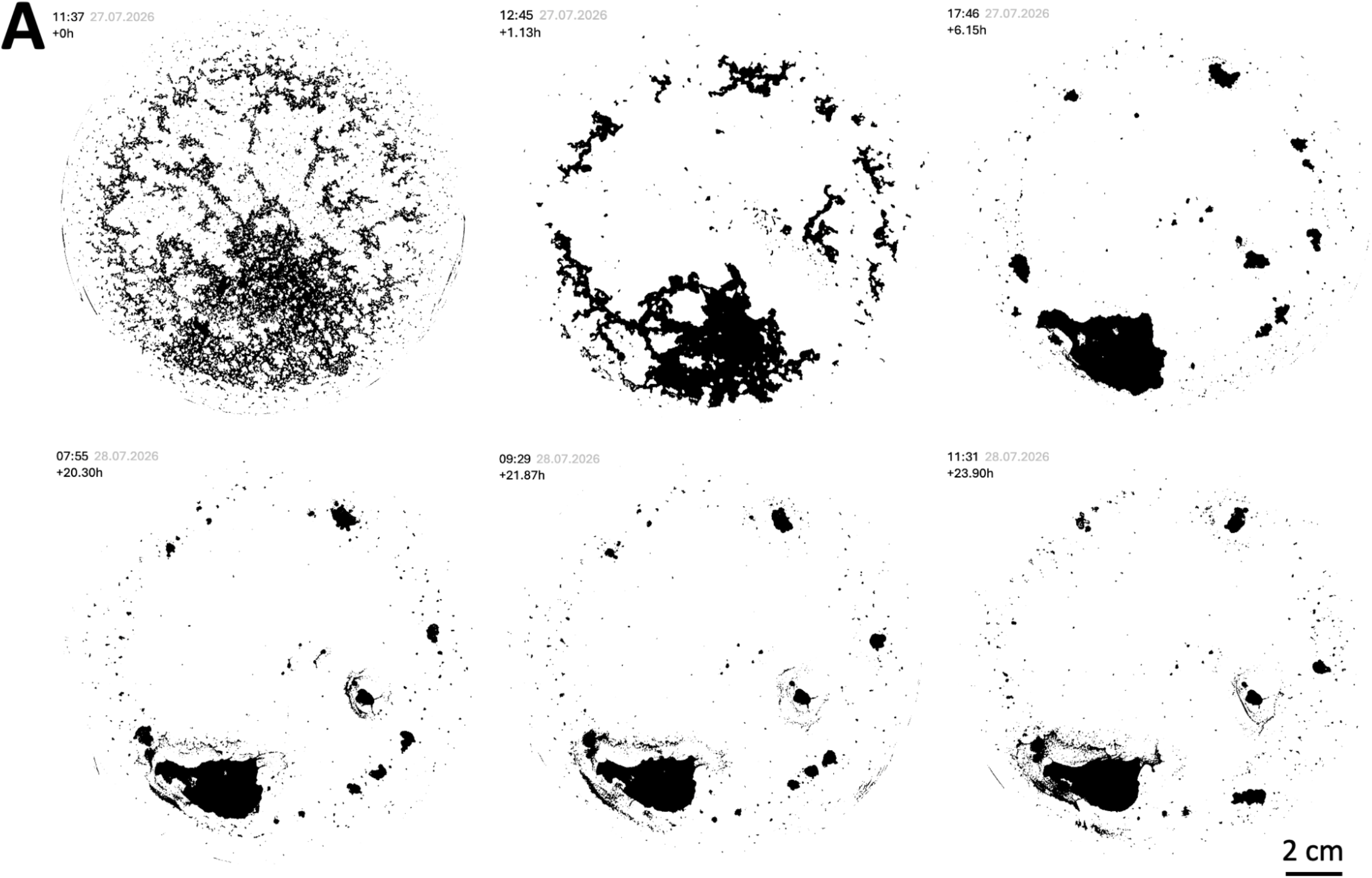
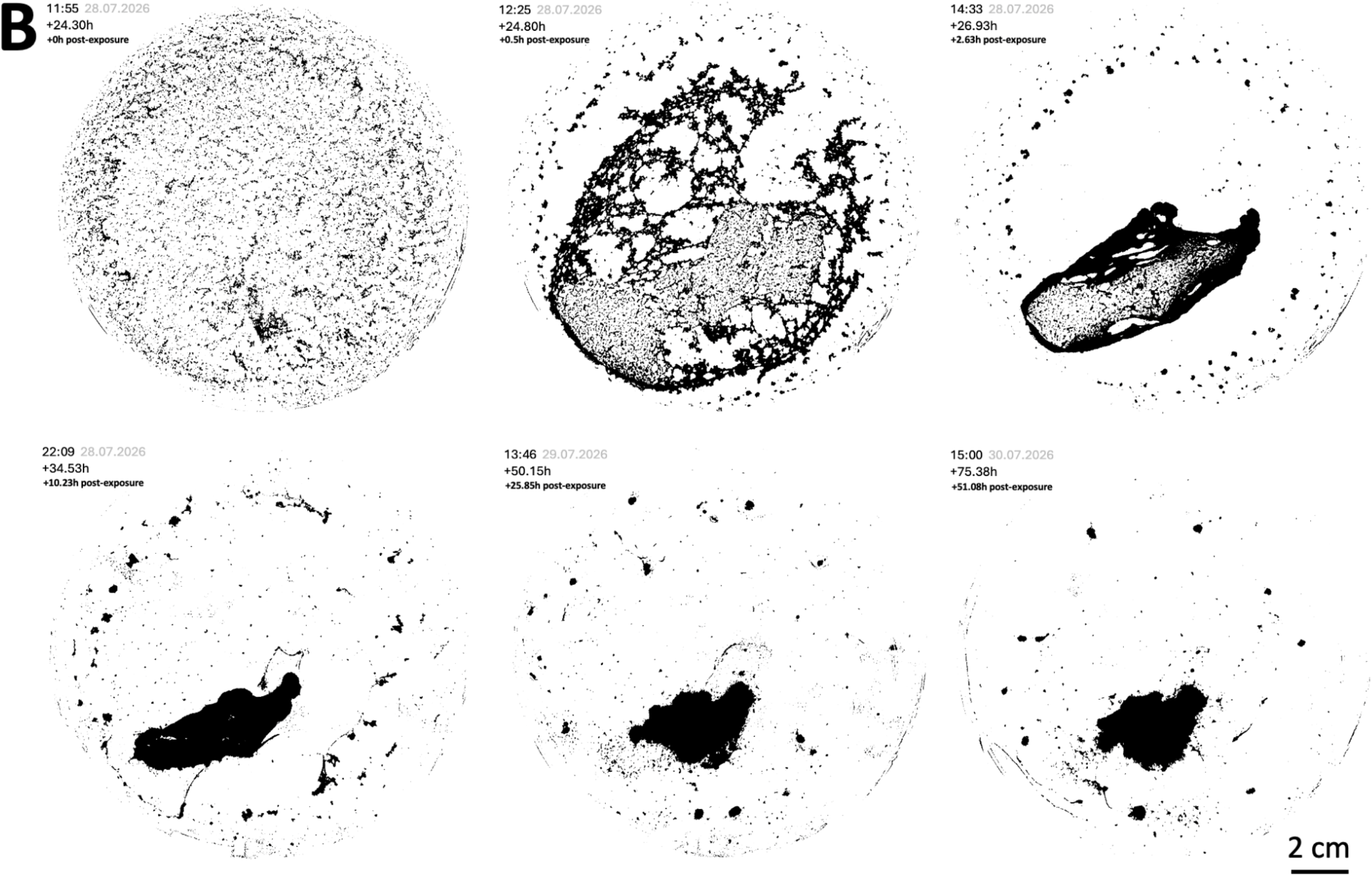
Temporal dynamics and structural organization of diatom aggregation during and after H_2_O_2_ exposure. Representative images of an H_2_O_2_-exposed natural microphytobenthic assemblage monitored over 72 h following resuspension from sediment. Images were repeatedly acquired from the bottom of the same 3-L Erlenmeyer flask containing 1 L of suspension. **A**. Representative temporal sequences showing changes in aggregation patterns over the first 24 h following H_2_O_2_ addition (nominal initial concentration of 400 µmol L^-1^); **B**. Representative temporal sequences showing changes in aggregation patterns during the 48 h post-exposure period, following medium replacement with filtered site water and resuspension.

Following resuspension from sediment, the natural assemblage rapidly formed multicellular aggregates, with diatom motility observed throughout the 72 h experiment under both control condition and H_2_O_2_ exposure (Figs. 1, 2; image-sequence animations and endpoint motility observations from the experiment: https://doi.org/10.5281/zenodo.22212357; time-lapse video from an additional experiment: Zenodo https://doi.org/10.5281/zenodo.22211925). Under the control condition, the initial aggregative structures progressively densified during the first hours, before developing into a spatially complex network comprising distinct structural forms, including compact aggregates and mushroom-like protrusions, as well as filamentous connections and biofilm-like mats (Fig. 1). Rather than progressing monotonically towards larger or denser structures, this organization underwent continuous dynamic remodeling throughout the 72 h experiment (Fig. 1A– C). Image analysis supported this dynamic: individual aggregate number changed from 11,767 at the beginning of the experiment to 3,237 after 72 h, while surface coverage changed from 24.4% to 10.3%, with pronounced fluctuations in both metrics between these endpoints (Supplementary Fig. 1 and Supplementary Table 1). Experimental observations further suggested that restructuring was spatiotemporally heterogeneous, with aggregate-specific and time-dependent structural reorganization rather than synchronous changes across the assemblage, as illustrated by the rapid but spatially localized changes observed between 7:55 and 9:29 h (Fig. 1A).

The formation and continual remodeling of these structures are consistent with the interplay between motility, cell adhesion, and extracellular material (Wang et al., 2021). EPS are particularly plausible contributors because they mediate diatom attachment and motility and contribute to biofilm organization (Hubas et al., 2018; Wang et al., 2021; Demir-Yilmaz et al., 2023). Consistent with a potential role of EPS in the observed aggregation, optically dense regions were observed within aggregates (Fig. 1F), which could reflect local EPS accumulation, although EPS was not specifically quantified here. However, the relationship between EPS production and aggregation is not necessarily proportional and may depend on environmental conditions, as shown in the marine diatom *Cylindrotheca closterium* (Demir-Yilmaz et al., 2023). Removal of most EPS markedly reduced, but did not completely prevent, aggregation, suggesting that EPS contribute substantially to aggregate formation but are not the sole determinant, with cell-surface properties such as hydrophobicity potentially also contributing (Demir-Yilmaz et al., 2023). From the present observations alone, it remains difficult to determine whether aggregation represents a biologically regulated collective behaviour, analogous to the endogenously regulated and environmentally modulated vertical migration of epipelic diatoms (Coelho et al., 2011; Farré et al., 2020; Desparmet et al., 2026), or an emergent consequence of motile encounters and cell–cell–matrix interactions (Wang et al., 2021). However, the persistent remodeling, spatiotemporally heterogeneous dynamics, and, as described below, alteration of aggregation dynamics under H_2_O_2_ exposure support the interpretation of a behaviour more complex than the passive accumulation of cells.

The aggregation dynamics observed under H_2_O_2_ exposure appeared to differ markedly from those observed in the control (Fig. 2A and Supplementary Fig. 1). After H_2_O_2_ addition, only 1,984 individual aggregates were detected, nearly sixfold fewer than in the corresponding control, while aggregate surface coverage remained comparable to the control (21.1% vs. 24.4%, respectively). After 24 h of exposure, aggregate number had decreased further to 1,182, more than eightfold fewer than in the corresponding control, while surface coverage declined to 6.0%, compared with 17.1% in the control (Supplementary Fig. 1 and Supplementary Table 1). In addition to these quantitative differences, the temporal pattern of aggregation was visually distinct: whereas aggregate number and surface coverage fluctuated markedly in the control, both metrics showed substantially less temporal variation during the H_2_O_2_ exposure period (Supplementary Fig. 1). These differences under H_2_O_2_ exposure were accompanied by a visibly altered structural organization, characterized by a dominant aggregate showing comparatively limited remodeling, while filamentous connections and biofilm-like mats were only sparsely observed and remained poorly developed compared with the control. During the subsequent 48 h post-exposure period, aggregate number and coverage remained comparatively weakly variable and ended at 841 aggregates and 6.2% coverage, without returning to the temporal dynamics observed in the control (Fig. 2B, Supplementary Fig. 1 and Supplementary Table 1). Nevertheless, motility remained visually apparent in H_2_O_2_-exposed diatoms at the end of the 72 h experiment (see https://doi.org/10.5281/zenodo.22212357), suggesting that the persistent difference in aggregation dynamics was not attributable to a loss of motility and may suggest a delayed relaxation of H_2_O_2_-responsive pathways and/or dependence of aggregation behaviour on the post-exposure physiological state of the cells.

Hydrogen peroxide is a well-established secondary messenger involved in stress responses, motility, and multiple signaling pathways in microalgae and plants, making the distinct aggregation dynamics observed here consistent with a biologically responsive component of aggregation behaviour (Waszczak et al., 2018; Morris et al., 2022; Desparmet et al., 2026). Under abiotic stress, aggregation has been reported as an acclimatory response in *Chlamydomonas reinhardtii*, in which aggregation increases with stress intensity and can enhance cellular protection (De Carpentier et al., 2022; Lambert et al., 2026). These observations raise the possibility that, in our experiment, the dense packing of diatoms within an EPS-rich biofilm matrix provided a locally buffered microenvironment following resuspension and exposure to exogenous H_2_O_2_ (Dang & Lovell, 2016; Flemming et al., 2016; Hubas et al., 2018). However, neither a protective effect of aggregation nor the underlying mechanism was directly tested here. Finally, regulation of aggregation in epipelic diatoms could involve extracellular secretions mediating intercellular perception, although the involvement of specific signaling molecules remains undemonstrated (Wang et al., 2021; De Carpentier et al., 2022; Yang et al., 2023; Lambert et al., 2026).

Overall, these observations identify rapid and continuously remodeled aggregation in a resuspended natural epipelic diatom assemblage and suggest that its temporal dynamics differed during and after H_2_O_2_ exposure. Although the experimental design did not isolate the effects of sediment removal and associated physical changes, nor specifically assess the contribution of the retained meiofauna, the distinct responses observed in the control and under H_2_O_2_ exposure suggest that aggregation dynamics are not temporally invariant and may respond to oxidative perturbation. Together with previous evidence of stress-induced aggregation and extracellular signaling in microalgae, these results raise the possibility that aggregation in epipelic diatoms includes an environmentally responsive, biologically regulated component. By promoting local buffering and biofilm organization, such regulation could enhance resilience to fluctuating intertidal conditions, although its adaptive functions remain to be tested. Distinguishing dynamics emerging from motile encounters and cell– cell–matrix interactions from those driven by regulated cellular responses now provides a testable framework for investigating collective behaviour and, ultimately, for better understanding the ecological success of motile diatoms (Nakov et al., 2018; Serôdio & Lavaud, 2020).

## Acknowledgements

We thank Hugo Le Gac for his assistance in the laboratory with the production of the time-lapse video of diatom aggregation, available here: https://doi.org/10.5281/zenodo.22211925

## Author contributions

**AD** (Conceptualization, Methodology, Investigation, Data curation, Formal analysis, Visualization, Writing - original draft); and **CH** (Supervision, Formal analysis, Validation, Writing - review & editing).

## Conflicts of interest

The authors declare no conflict of interest.

## Funding

This work was funded by the Institut de l’Océan from the Sorbonne University alliance and the Muséum national d’Histoire naturelle.

## Data availability

All scripts and data presented in this article and used to generate the figures and perform the analyses are available on GitHub (https://github.com/adesparmet/Epipelic-diatoms-aggregation).

## Supplementary data

**Figure S1.**
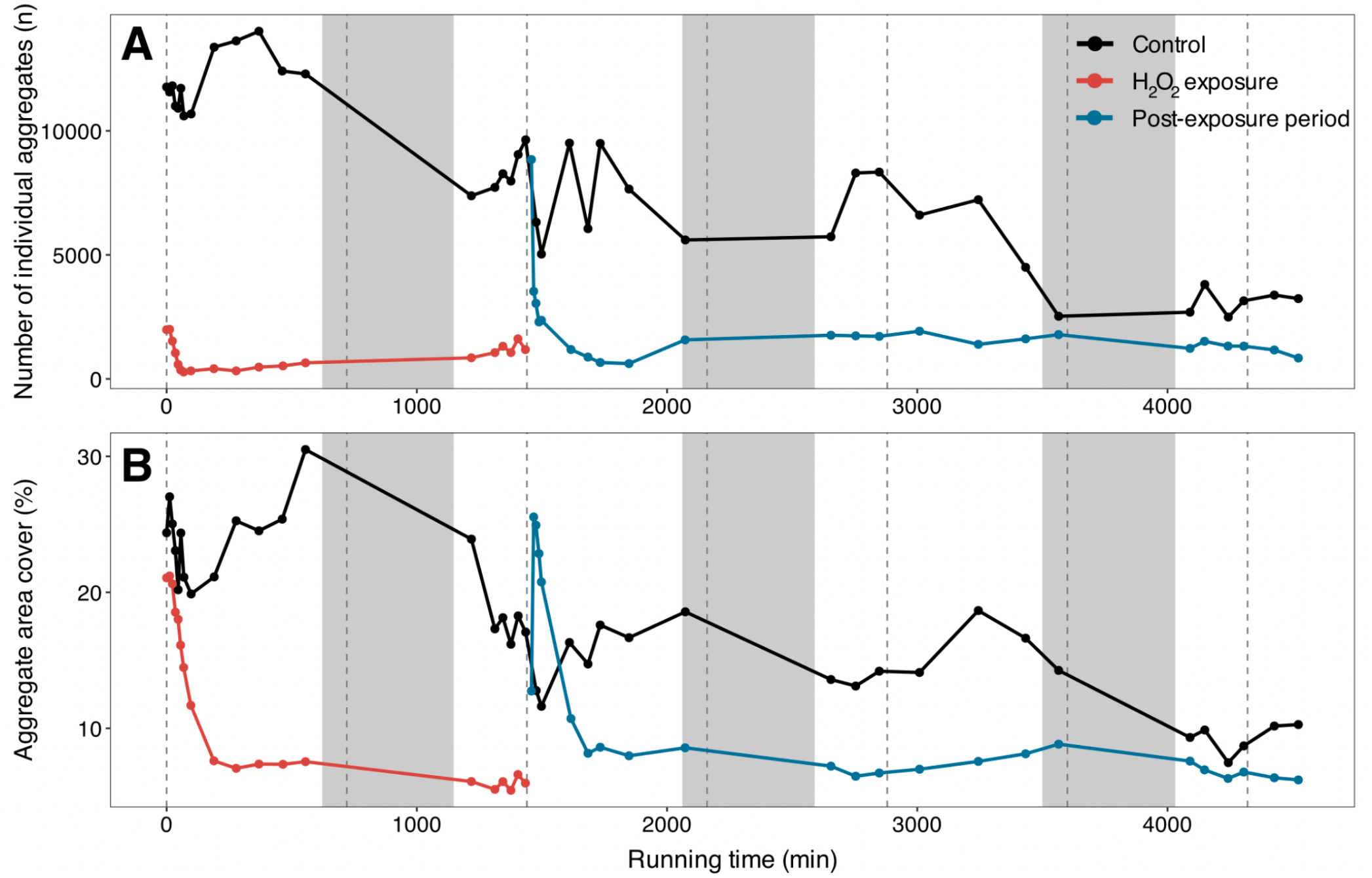
Temporal dynamics of diatom aggregate formation under control condition and H_2_O_2_ perturbation. Image-based analysis (Supplementary Fig. 2) of **A**. the number of individual aggregates (n, *n* = 1) and **B**. the area covered by aggregates relative to the available colonizable surface (%, *n* = 1), over the course of the experiment, under the control condition (black), H_2_O_2_ exposure (red), and during post-exposure period (blue). Solid lines connect successive observations; points indicate individual image-derived measurements. Grey shading highlights the local natural nighttime period. Dashed vertical lines indicate 12-h intervals (noon and midnight). Detailed raw data are provided in Supplementary Table 1. *Images were analysed using the EBImage package. Aggregate number was calculated as the number of individually detected objects, whereas area cover was calculated as the summed aggregate area divided by the total area available for colonization on the Erlenmeyer bottom × 100. Experimental time is expressed as elapsed time in minutes from the beginning of the experiment. Note that, in panel B, the first post-exposure measurement (blue) was performed shortly after resuspension, before complete diatom resettlement, resulting in an initial increase in aggregate area cover as the remaining diatoms resettled. This measurement was therefore not considered as the initial reference time point for the post-exposure condition*.

**Figure S2.**
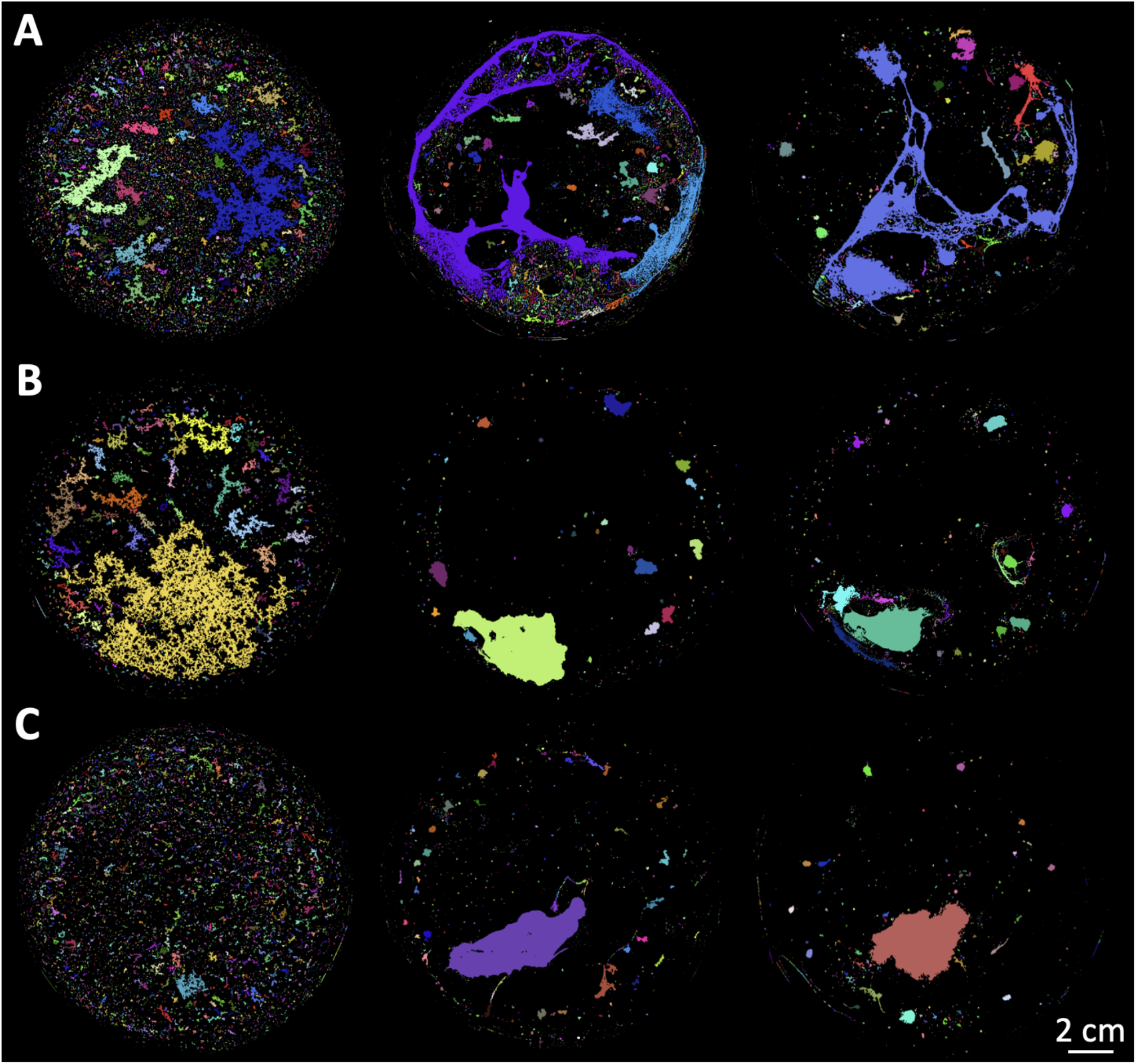
Representative images illustrating the image data acquisition procedure. Examples of processed images generated using the EBImage package, corresponding to representative images presented in Figs. 1 and 2 and used to derive the aggregate number and area cover metrics. Colours assigned to individual detected objects are random and are therefore intended for visualization only; colour identities are not comparable between images. **A**. Images of aggregation under the control condition; **B**. Images of aggregation during H_2_O_2_ exposure; **C**. Images of aggregation during the post-exposure period.

**Table s1.**
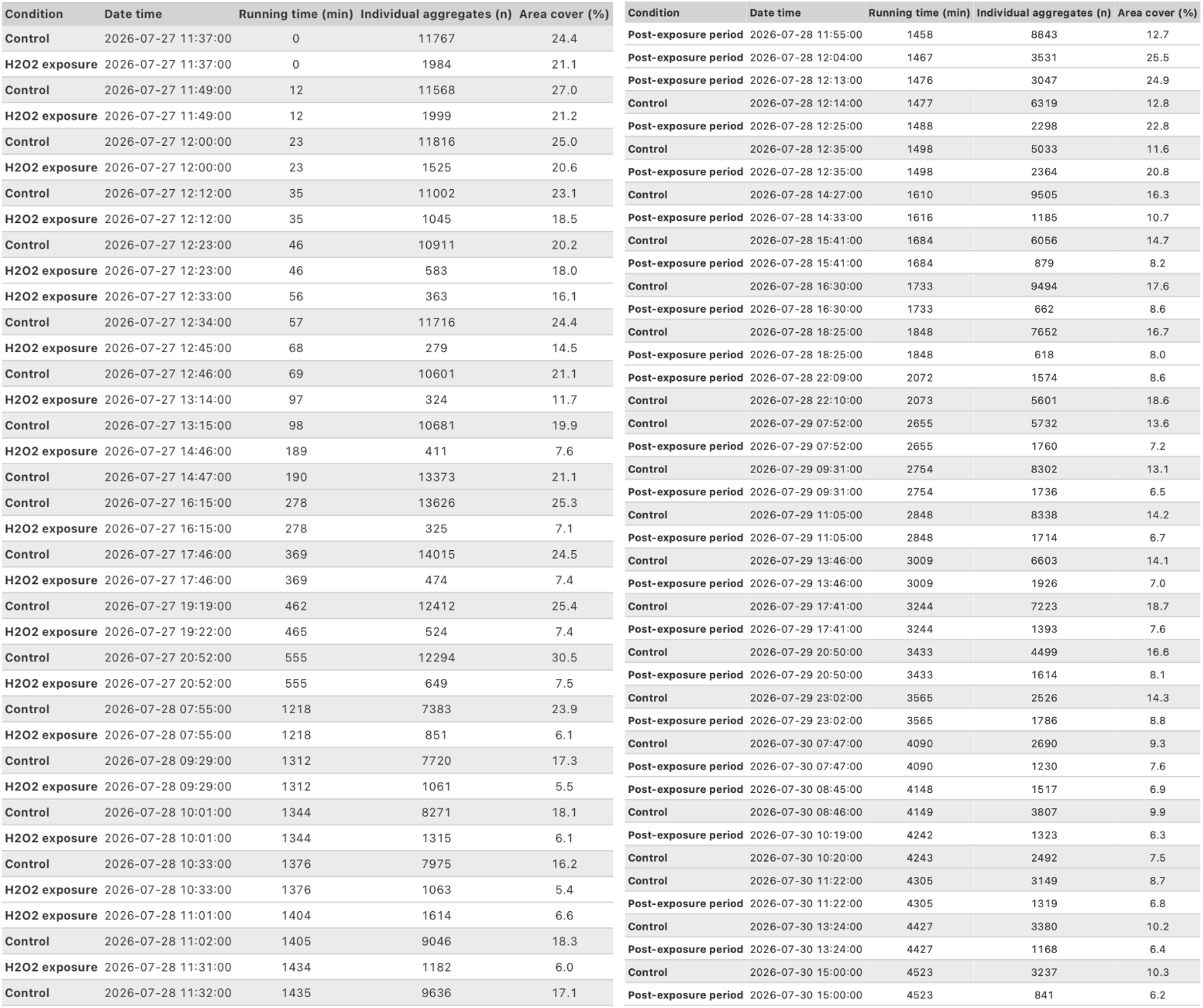
Image-based quantification of diatom aggregate dynamics over time. Temporal records of the number of individual aggregates and the area covered by aggregates, under the control condition, during H_2_O_2_ exposure, and during the subsequent post-exposure period. Each row corresponds to an individual image acquisition. “Condition” indicates the experimental condition; “Date time” corresponds to the date and time of image acquisition; “Running time (min)” indicates elapsed experimental time; “Individual aggregates (n)” corresponds to the number of individual aggregates detected in the image (*n* = 1); and “Area cover (%)” represents the summed aggregate area relative to the total surface available for colonization on the Erlenmeyer bottom (*n* = 1). Control measurements are highlighted in grey. *Image analysis was performed using the EBImage package*.

## Notes

https://github.com/adesparmet/Epipelic-diatoms-aggregation

### Competing Interest Statement

The authors have declared no competing interest.

